## Supplementary Data 7 for "Metabolic modeling reveals a multi-level deregulation of host-microbiome metabolic networks in IBD"

### Supportive information

#### Description of the general network topology

We based the network reconstruction on significantly associated reactions with the tested conditions in IBD patients, separated by tissue (blood and gut biopsy). Reactions formed the edges, while their substrates were displayed as nodes. The network topology was characterized using the log of edge centrality as a measure of distance between the nodes (Supplementary Fig. 3).

The analysis revealed two major points: Firstly, the biopsy network topology was largely unstructured, comprising a single large network without distinct subnetworks (Supplementary Fig. 4A), except for a small cluster involving maltose metabolism (Supplementary Fig. 4B). In contrast, the blood network had a central main subnetwork and three additional subnetworks. Each of these subnetworks contained at least one central metabolite, indicating their respective function – one involved D-galactose and carbohydrate metabolism. Another was associated with lipid metabolism (containing cholesterol, cholesterol-ester, phosphatidylcholine, and phosphatidylethanolamine). The last subnetwork contained reactions for amino acid metabolism indicated by the presence of arginine, tryptophan, lysine and histidine in the network (Supplementary Fig. 4B). Secondly, the hub metabolites (Supplementary Fig. 4C) were centrally located within the networks/subnetworks, underlining their importance in the inflammation-induced changes in host metabolic activity. Notably, the cofactors NAD/NADP(H) were positioned at the center of both networks, emphasizing their significance in inflammation-related host metabolism.

To understand the positioning of hub metabolites (compare Fig. 3B) as well as microbial metabolites, and those derived from metabolomic data within the network topologies, we highlighted the nodes and their connecting reactions (Supplementary Fig. 4C, D). Next, we aimed to comprehend how these metabolites are interconnected with each other as a proxy for the interdependence between microbial and host metabolism, as well as blood metabolomics.

#### Detailed description of modeling results for remission and responding patients

For the microbial changes, we observed reduced NAD (de-)phosphorylation (Supplementary Fig. 6B) and increased production of 3-dehydrocholate (Supplementary Fig. 6D) in responding patients. The former fits to the observation of more de-novo NAD synthesis during inflammation, while the latter indicates more active de-conjugation of bile acids. Surprisingly, we observed a reduced activity of the mixed acid fermentation pathway for responders (Supplementary Fig. 6B), which predominantly produces lactate, acetate, and formate, but not butyrate or propionate. This correlates to the increased exchange of lactate, decreased microbial exchange of propionate, and reduced host-relevant butyrate production during inflammation (Supplementary Fig. 6B, Fig. 2C, D).

For remission, we observed an increase in pyrimidine salvage and nucleotide synthesis (Supplementary Fig. 6F – both pathways necessary for nucleotide and NAD synthesis (reduced during inflammation, Fig. 2B). Nicotinamide was also less exchanged in responder microbiomes and more available for the host (Supplementary Fig. 6G, H). For amino acids, we found reduced synthesis of lysine, leucine, and arginine (Supplementary Fig. 6F), resulting in less arginine (and glutamine) cross-feeding between bacteria (Supplementary Fig. 6G). Concurrently, we observed increased consumption of dietary proline with treatment response (Supplementary Fig. 6H). This contrasts with the increased host availability of leucine, asparagine, and proline during inflammation (Fig. 2D). Hence, competition for proline seems crucial for healthy microbiome-host interactions.

Further, we observed increased production of monosaccharides (xylitol, glucose) and their fermentation products (malate), while complex carbohydrates (starch, malto-11-ose) were less available to the host (Supplementary Fig. 6H). This suggests an enhanced degradation of resistant carbohydrates and their usage in fermentation processes in responder microbiomes. Homocysteine-methionine-cysteine interconversion was reduced in the response phenotype (Supplementary Fig. 6F), indicating an increased availability of these compounds, including homocysteine, which was identified as central to the metabolic changes during inflammation in the host (Fig. 7G, H). Additionally, we observed a decrease in synthesis of fatty acids, teichoic acid, phosphatidylethanolamines, and farnesol (Supplementary Fig. 6F) which could result in reduced usage of SCFAs in these pathways and increased availability for the host. Furthermore, we identified increased β-alanine production as positively associated with remission. Beta-alanine is produced from propionate and is used to produce coenzyme A, thereby important for lipid metabolism (Supplementary Fig. 6F). This relates back to the inflammation-induced changes in the microbial metabolism, where we observed decreased propionate exchange among bacteria, thus less usage of propionate in their own metabolic processes. We also found decreased synthesis of lipoteichoic acid and arachidonoylglycerol, indicating a reduced demand for coenzyme A (Fig. 2B, C).

For the changes in host metabolism, we found increased amino acid metabolism (arginine, proline, glycine, serine, alanine, threonine) and increased glutathione metabolism in patients responding to treatment (Supplementary Fig. 7C), reversing some effects observed during inflammation. In patients undergoing remission, we observed increased sphingolipid metabolism in biopsies, which were reduced during active disease in blood (Supplementary Fig. 7G). Hence, the gut seems to compensate for some of the changes observed in blood during remission.

When analyzing metabolite enrichment in the significantly associated reactions with response and remission, we found no consistent patterns between blood and gut tissue (Supplementary Fig. 7D, H). Notably, for response, the enrichment of NAD(H) in biopsies and coenzyme A in blood was driven by more upregulated reactions in responsive patients, contrasting with the downregulation of these metabolites during inflammation (Supplementary Fig. 7D).

In the analysis of the metabolomics results associated with response and remission, we observed an increase of serum levels of lysophosphatidylcholines, phosphatidylcholines, and choline (response only) (Supplementary Fig. 8A, D), contrary to the decreased levels observed during inflammation (Fig. 4A). This further indicates increased availability of choline for recycling homocysteine to methionine in patients responding to the treatment (Fig. 6G, H). Relatedly, we found increased levels of cystine (cys-S-S-cys) in the same analysis, another indicator of sufficient homocysteine levels and a substrate for glutathione production (Supplementary Fig. 8A, Fig. 6E-H). In addition, we identified higher levels of kynurenine and serotonin in treatment-responsive patients (Supplementary Fig. 8A), while we found no association with tryptophan, indicating sufficient levels of tryptophan to fuel de-novo NAD synthesis (Fig. 6A-D) and even production of serotonin from it.

Finally, we repeated the network reconstruction to find interconnections between microbial, host, and blood metabolomics metabolites for patients in response and remission. Due to the fewer reactions identified in the host metabolism under these conditions, we recovered smaller networks for all conditions (Supplementary Fig. 9, Supplementary Data 3, Supplementary Data 4, Supplementary Data 5, Supplementary Data 6). Especially for remitting patients, we recovered networks too small to establish any direct interactions between the different metabolites (Supplementary Fig. 9E-H). For response, we again found significantly shorter paths between center, microbial, and metabolomic metabolites than between all other metabolites in the network (Supplementary Fig. 10C). Nonetheless, in both tissues, we found similar connections between the choline to phosphatidylcholines (Cen-Met) and phosphatidylcholines to homocysteine (Mic-Met) as in the inflammation network (Supplementary Fig. 10B, D).
